# Epigenetic models predict age and aging in plains zebras and other equids

**DOI:** 10.1101/2021.03.29.437607

**Authors:** Brenda Larison, Gabriela M. Pinho, Amin Hagani, Joseph A. Zoller, Caesar Z. Li, Carrie J. Finno, Colin Farrell, Christopher B. Kaelin, Gregory S. Barsh, Bernard Wooding, Todd R. Robeck, Dewey Maddox, Matteo Pellegrini, Steve Horvath

## Abstract

Five of the seven extant wild species of the genus *Equus* are species of significant conservation concern. Effective conservation and management of such threatened wildlife populations depends on the ability to estimate demographic trends and population viability and therefore requires accurate assessment of age structure. However, reliably aging wildlife is challenging as many methods are highly invasive, inaccurate, or both. Epigenetic aging models, which estimate individual age with high accuracy based on genomic methylation patterns, are promising developments in this regard. Importantly, epigenetic aging models developed for one species can potentially predict age with high accuracy in sister taxa. Using blood and biopsy samples from known age plains zebras (*Equus quagga*), we developed epigenetic clocks (ECs) to predict chronological age, and epigenetic pacemaker (EPM) models to predict biological age. We tested the ability of our blood-based EC to predict ages of Grevy’s zebras, Somali asses and domestic horses, from blood samples. Because our samples came from a population with a complex pedigree, we also leveraged information from a previous sequencing effort to measure the association between levels of inbreeding (F and ROH) and the age acceleration as measured by DNA methylation. The resulting models describe the trajectory of epigenetic aging in plains zebras and accurately predict the ages of plains zebras and other equids. We found moderate support for a slight acceleration of aging with increased inbreeding.

## Introduction

Effective management of threatened species relies on the ability to estimate demographics trends, which rely, in turn, on accurate information about age distributions within populations [1]. Age distributions are shaped by growth rates and can reflect past and current environmental and anthropogenic perturbances [2, 3] and can also be used to predict future population growth [4]. However, age is challenging to quantify in wild animals. Age estimation typically requires either invasive approaches that may not be feasible in live animals or investment in long term field studies [2, 4, 5]. Another problem is the limited accuracy of some methods, which may negatively impact conservation efforts [2]. The challenges and importance of obtaining accurate age information has motivated efforts to develop an accurate and non-invasive approach to aging animals [4, 6].

Epigenetic clocks (ECs) are a promising development for the aging of wild animals and have the potential to make significant contributions to wildlife conservation and population biology [4, 6]. Such clocks are highly accurate and there is a rich literature focusing on ECs in humans [7–9] and mice [10–12], however the availability of ECs for other species, while increasing, is still limited [6]. A critical limitation to developing epigenetic aging models for wildlife is that these models need to be trained on samples from individuals of known age, therefore populations of non-model organisms with known-age individuals are of extreme importance [6, 13].

Besides their accuracy, four other features of epigenetic aging models should make them attractive for wildlife managers: First, they can be developed from different tissue types [6, 14]. Second, epigenetic clocks can be created based on very few genomic sites. The most accurate clocks for humans involve only a few hundred CpG sites [14, 15], and far fewer CpG sites have been used to build epigenetic clocks in some wild vertebrates [3, 16, 17]. Third, an accurate clock can be developed using relatively few individuals of known age [3, 6, 17]. Finally, epigenetic clocks developed for one species have been shown to accurately predict age in closely related taxa (e.g., Humans and chimps; [14]), so epigenetic clocks can be developed for a less threatened species with the intent of using them in threatened sister species.

When individual chronological ages are known, it is possible to estimate how fast individuals are aging compared to others by the discrepancy between epigenetic age and chronological age, dubbed age acceleration. A positive age acceleration indicates that an individual is biologically older than expected based on its chronological age. Age acceleration is predictive of all-cause mortality in humans [18–22], which suggests that ECs can be a powerful approach to study the impact of different factors on biological aging. Age-acceleration has also been associated with stress and adversity [23–25], elevated glucocorticoids [26, 27], and inbreeding [28–30], all of which are relevant for managing wild populations.

The extraordinary accuracy of ECs stems from their utilization of sites that maximize a linear relationship between epigenetic age and chronological age. However, epigenetic age changes in a non-linear fashion throughout individuals’ lifetimes in several species, with accelerated changes in early life and slower changes once individuals reach adulthood [14, 31]□. In order to model these nonlinear changes in methylation levels with chronological age, the epigenetic pacemaker (EPM) has recently been developed [31–33]. The EPM estimates epigenetic age by maximizing the similarity between estimated and observed methylation levels, and therefore does not make any assumptions about linearity but rather identifies the shape of the relationship between age and methylation directly from the data. In this sense, the EPM is potentially more associated with biological, rather than chronological aging [31], which is particularly useful for investigating how environmental and life-history factors influence the aging process. For instance, it has recently been used to investigate aging in the yellow-bellied marmot, a hibernating species, showing that epigenetic aging slows during hibernation [34].

Here, we develop the first epigenetic models for a wild equid (plains zebras, *Equus quagga*) using both blood and remote biopsy samples collected from known-age individuals of a captive-bred population. We test the ability of the blood-based EC to predict age in three closely related equid species. Using EC- and EPM-estimated predictions we examine whether inbreeding is associated with accelerated epigenetic aging in this population. The development of epigenetic aging models in a wild equid stands to have broad impact because the crown group of the genus *Equus*, comprises a closely related group of 6 extant species, 5 of which range from near threatened to critically endangered [35, 36]. Because the speciation events within *Equus* are more recent than between humans and chimps (< 2 MY vs. 6-8MYA; [37, 38] the models we develop in plains zebras are likely to have significant utility in other equids.

## Methods

### Samples

Both whole blood (96) and remote biopsy (24) samples were obtained from a captive population of zebras maintained in a semi-wild state by the Quagga Project [39] in the Western Cape of South Africa. The population was founded in 1989 with 19 wild individuals (9 from Etosha National Park in Namibia, 10 from the Kwazulu-Natal in South Africa). Since its inception, the population has undergone strong selection in an effort to reproduce the phenotype of the extinct quagga subspecies: no stripes on legs and hindquarters, and thinner and paler stripes in the head and barrel region. During sampling, individuals were uniquely identified by their stripe patterns and their ages were derived from studbook information about dates of birth, which were typically known within one month. One exception to this is a tissue sample from a founder that was sampled as a young mare and would have been at least 25 years old at sampling, but may have been slightly older. Remote biopsies were taken using an air-powered rifle affixed with a 1 mm wide by 20-25 mm deep biopsy dart and preserved in RNAlater (Qiagen). Blood samples were collected opportunistically during veterinarian visits, and preserved in EDTA tubes. Most samples were collected from different individuals, with the exception of two individuals that each were sampled twice some years apart. All samples were stored at −20 °C. After eliminating samples with low confidence for individual identity and age, we retained 76 blood samples and 20 biopsy samples, totaling 96 zebra samples (**Table 1**, Supplement 1).

**Table 1.** Description of the zebra data. We restrict the description to animals whose ages could be estimated with high confidence (90% or higher). Tissue type, N=Total number of samples/arrays. Number of females. Age: mean, minimum and maximum.

| Tissue | N | No.<br>Female | Mean<br>Age | Min.<br>Age | Max.<br>Age |
| --- | --- | --- | --- | --- | --- |
| Blood | 76 | 42 | 5.21 | 0.156 | 20.2 |
| Skin | 20 | 9 | 5.87 | 0.162 | 24.8 |

Three zebra data sets were analyzed in our epigenetic models: (1) only blood samples, (2) only biopsy samples, and (3) blood and biopsy samples combined. In addition, we evaluated the feasibility of applying the blood-based zebra clock to predict ages of other equids using known age domestic horses (*E. caballus*), and a small set of samples of known-age individuals from two closer relatives, Grevy’s zebras (*E. grevyi*) and Somali asses (*Equus africanus somaliensis*). The collection of 188 whole blood samples from domestic horses are described in detail in [40]. The Grevy’s zebra (n=5) and Somali wild ass (n=7), are samples from zoo-based animals that were opportunistically collected and banked during routine health exams and the DNA methylation profiles from these samples have been reported previously [41].

### Ethics approval

Plains zebra samples were collected under a protocol approved by the Research Safety and Animal Welfare Administration, University of California Los Angeles: ARC # 2009-090-31, originally approved in 2009.

### DNA methylation data

All DNA methylation data (plains zebra, horse, Somali wild ass, Grevy’s zebra) were generated using a custom methylation array (HorvathMammalMethylChip40) [42]. The array contains 36 thousand probes, 31,836 of which mapped uniquely to the horse genome [43, 44]. Methylation values from zebra blood and biopsy tissue were each normalized separately using SeSaMe [45]. The same SeSaMe normalization method was performed for all tissue samples from all species (plains zebra, horse, Somali wild ass, and Grevy's zebra). Unsupervised hierarchical clustering revealed that the plains zebra samples clustered by tissue (Supplement 3, Figure S1).

### Epigenetic Aging models

We studied epigenetic aging in plains zebras using both epigenetic clock (EC) [14, 46, 47] and epigenetic pacemaker (EPM) models [31–33]. The ECs were developed by fitting a generalized linear model with elastic-net penalization (alpha=0.5) using leave-one-out (LOO) cross-validation in glmnet v.4.0-2 in R [48, 49]. To improve EC fit [14] we square root transformed chronological age prior to fitting the models. We report the coefficients, intercepts and lambdas for the final models, trained on all samples, in Supplement 2. The data from other equids (domestic horse, Grevy’s zebra, Somali wild ass) were treated as independent data, i.e. these data were not used in any way to build the zebra clocks.

To construct the EPMs we used sites in which methylation levels were highly correlated with individual chronological age. The Pearson correlation (*r*) thresholds for entry into the model were absolute values of 0.75 for blood and biopsy, and 0.6 for the combined EPM. The threshold used to select the sites for input into the combined EPM was lower because only one CpG site had *r* higher than 0.75. Epigenetic states were estimated using a leave-one-out cross validation with EpigeneticPacemaker 0.0.3 [32] in Python 3.7.4 [50]. All Pearson coefficients of methylation with plains zebra chronological age, and the rate and intercept values per site for blood, biopsy and combined data sets are provided in Supplements 2 and 4.

### Association of Inbreeding with Biological Aging

We used predictions from blood-based EC and EPM to assess whether inbreeding is associated with age acceleration (predicted age – age) in this population. Sex did not have a significant effect on age acceleration, so was not used in the regressions. The genotypes used in the estimation of F and ROH were derived from two sources: RADseq data (39 individuals) and genotypes imputed for 26 individuals at the same RADseq sites from whole genome low coverage sequence data. Details of library preparation and sequencing can be found in Supplement 3. Imputations were performed in GLIMPSE [51] with a likelihood threshold of 0.1 for accepting homozygote calls and of 0.9 for accepting heterozygote calls. We combined the RADseq reference panel to the imputed samples. This resulted in a sample size of 66 individuals spanning generations 1-6 of the Quagga Project. We imposed moderate pruning as recommended by [52] using a variance inflation factor (vif) of 2 in PLINK [53] leaving 227882 SNPs for estimating inbreeding coefficients. We estimated F and ROH in PLINK. F is estimated in PLINK using methods of moments. We used the default settings for ROH with the exception that no more than 2 missing SNPs per window were allowed. This results in a minimum ROH size of 1 MB. ROH was converted to the inbreeding coefficient FROH by dividing total MB in ROH by 3 GB which is the approximate size of the horse genome. We regressed inbreeding coefficients on age-acceleration using nonparametric Theil-Sen Siegel regressions [54] with the package mblm [55] in R.

### Epigenome Wide Association Studies (EWAS) of Age and Functional Enrichment

EWAS was performed in each tissue separately using the R function “standardScreeningNumericTrait” from the “WGCNA” R package [56]. Next the results were combined across tissues using Stouffer’s meta-analysis method [57]. We estimated the position of each significant CpG site in relation to the closest transcriptional start site. From each EWAS (blood, skin, and meta-analysis) we retained the 500 CpGs with the most positive z-scores and the 500 with the most negative z-scores. These CpGs were used as the input for GREAT analysis software [58]. The background was human Hg19 genome, limited to 31,836 CpG sites that could be mapped the horse genome. The options in the analysis included “Basal plus extension” and a maximum of 50 kb flanking window for the CpGs coordinates.

## Results

### Epigenetic Aging models

The blood EC (Pearson’s *r* = 0.96, median absolute error (MAE) = 0.56 years, **Figure 1A**) and the combined tissue EC (*r* = 0.89, MAE = 0.62, **Figure 1C**) predict age more accurately than the biopsy EC (*r* = 0.62, MAE = 1.79 years, **Figure 1B**). The blood EC used 70 CpG sites, the biopsy clock 31 sites, and the combined clock used 99 sites. The biopsy EC has no CpG sites in common with the blood and combined ECs. The blood EC and combined tissue EC share only two CpG sites.

**Figure 1.**
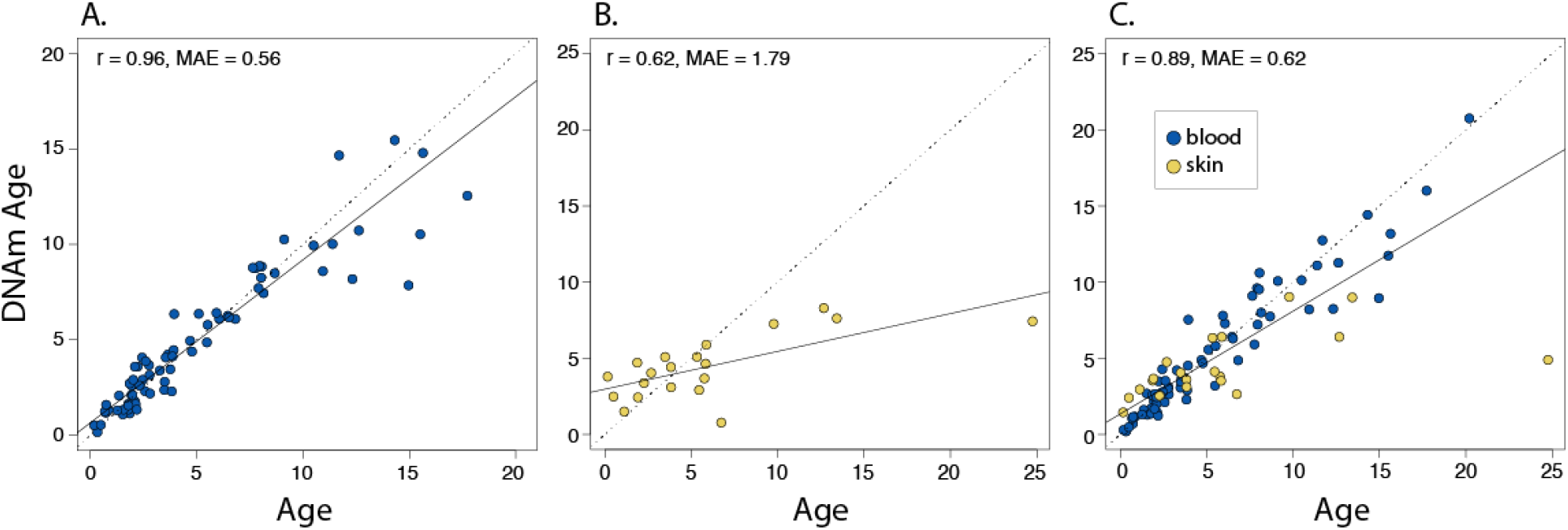
Epigenetic clocks for plains zebras. We developed 3 epigenetic clocks for zebras using square-root transformed ages: A) epigenetic clock for blood samples, B) epigenetic clock for biopsy samples, and C) combined tissue clock for both sample types. Leave-one-sample-out (LOO) estimate of DNA methylation age (y-axis, in units of years) versus chronological age (x-axis). In A-C the linear regression of epigenetic age is indicated by a solid line while the diagonal line (y=x) is depicted by a dashed line.

Cross-species predictive ability was high (**Figure 2)**. The zebra-blood EC predicted horse age with high accuracy (*r* = 0.93, MAE = 1.82). The error when predicting the ages of horses younger than 15 years of age is lower (MAE 1.15) than when predicting the ages of older horses (MAE = 3.97). While prediction errors for Grevy’s zebra and Somali wild ass ages were even lower (MAE of 1.08 and 1.15 respectively), we caution the reader that we had only limited sample sizes from Grevy’s zebra (n=5) and Somali wild ass (n=7).

**Figure 2.**
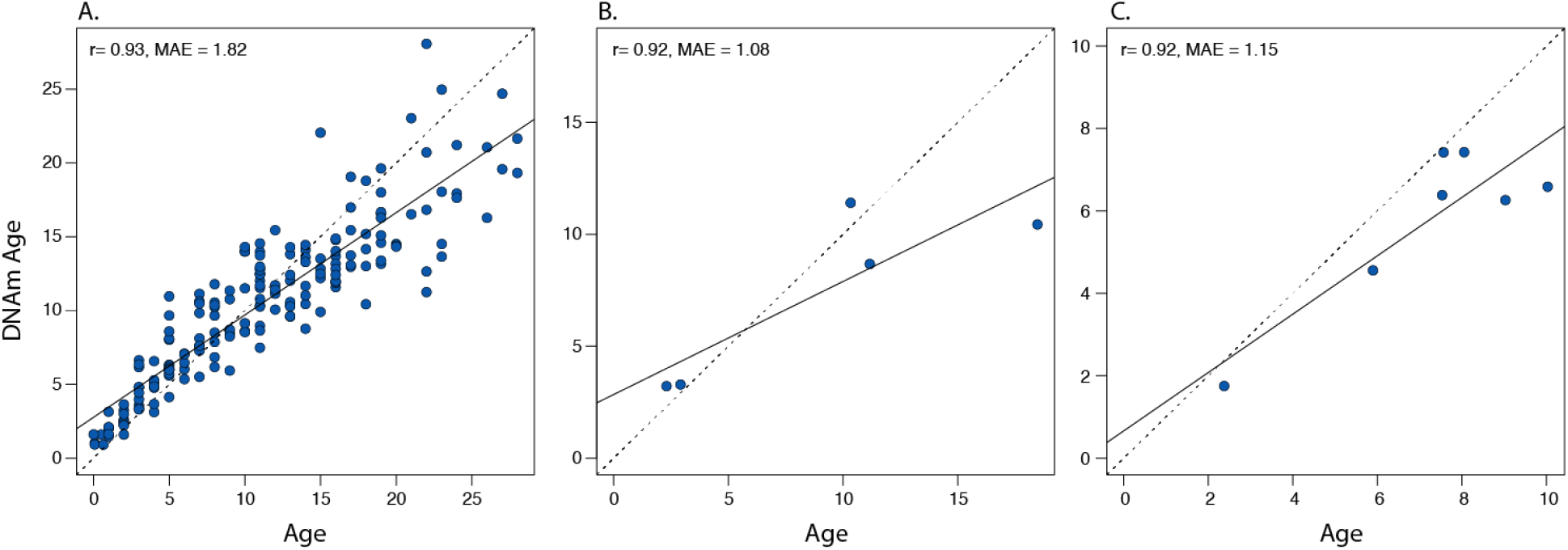
Zebra EC applied to samples from other equids. Blood samples from three sister taxa were used to test the ability of the plains zebra blood clock to predict chronological age in other equids: A) domestic horse N=188, B) Grevy’s zebra N=5, C) Somali wild ass N=7.

The blood, biopsy and combined-tissue EPMs included 391, 242 and 248 sites CpG sites, respectively. Epigenetic state was strongly correlated with chronological age in both tissue types: blood (*r* = 0.97, **Figure 3A**) and biopsy (*r* = 0.95, **Figure 3B**). EPMs based on the two tissues used largely distinct sets of CpG sites, sharing only 40 sites. Despite retaining an overall strong correlation in the combined EPM (*r* = 0.96), the difference in the two sample types is apparent (**Figure 3C**). The combined tissue pacemaker shared 138 sites with blood and only 34 with biopsies. Details of the CpGs selected by each model are in Supplement 2.

**Figure 3.**
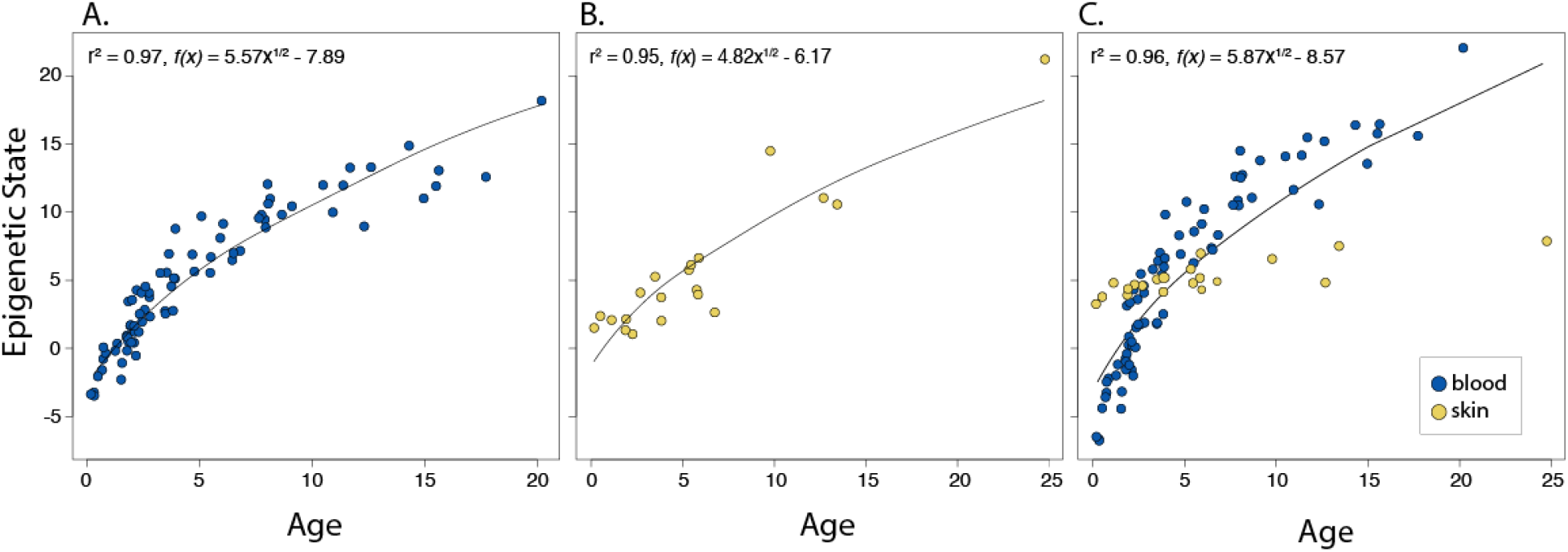
Epigenetic pacemaker (EPM) models for plains zebras. Epigenetic states of plains zebras predicted from the epigenetic pacemaker model using A) blood (391 CpG sites) MAE of translated ages 1.27, B) remote biopsy tissue (242 CpG sites) MAE of translated ages 1.28, and C) both sample types combined (248 CpG sites) MAE of translated ages 1.77. Predictions are based on 76 blood samples and 20 biopsy samples.

### Association of Measures of Inbreeding with Biological Aging

F ranged from −0.53 to 0.31, F_ROH_ ranged from 0 to 0.3. F and F_ROH_ were strongly correlated (r= 0.82, t=11.42, *P*= 2.2e-16). We detected a significant positive association of F and age acceleration based on both epigenetic age and state (Supplement 3, Tables S1 and S2). We also found a significant positive association of F_ROH_ with epigenetic age (EC) but not state (EPM). (Supplement 3, Tables S1 and S2).

### EWAS of chronological age in skin and blood of zebras

The current analysis is based on the 3,1836 probes in mammalian methylation array that could be aligned to specific loci adjacent to mammalian methylation array in horse genome. Since mammalian methylation array is based on the conserved stretches of DNA in all mammals, the horse annotation can be applied to the zebra data [42]. At a nominal p<10^−4^, a total of 9757 and 331 probes were related to age in blood (n = 76, age range 0.15-20.2 years) and skin (n = 20, age range 0.16-24.8 years) respectively (**Figure 4A**). The top age-related changes per tissue is as follows: blood, hypomethylation in *FANCL* upstream, *MAF* downstream, *ZNF608* upstream, and PBX3 intron; skin, hypermethylation in *PLCB1*, *NEUUROD1*, and *BARHL2* upstream. DNAm aging was distributed in all genic and intergenic regions relative to transcriptional start site (**Figure 4B**). Promoters and 5’UTR regions, which can be considered as expression regulatory regions, mainly gained methylation in both tissues. This observation paralleled a systematic positive correlation of CpG islands with age (**Figure 4C**).

**Figure 4.**
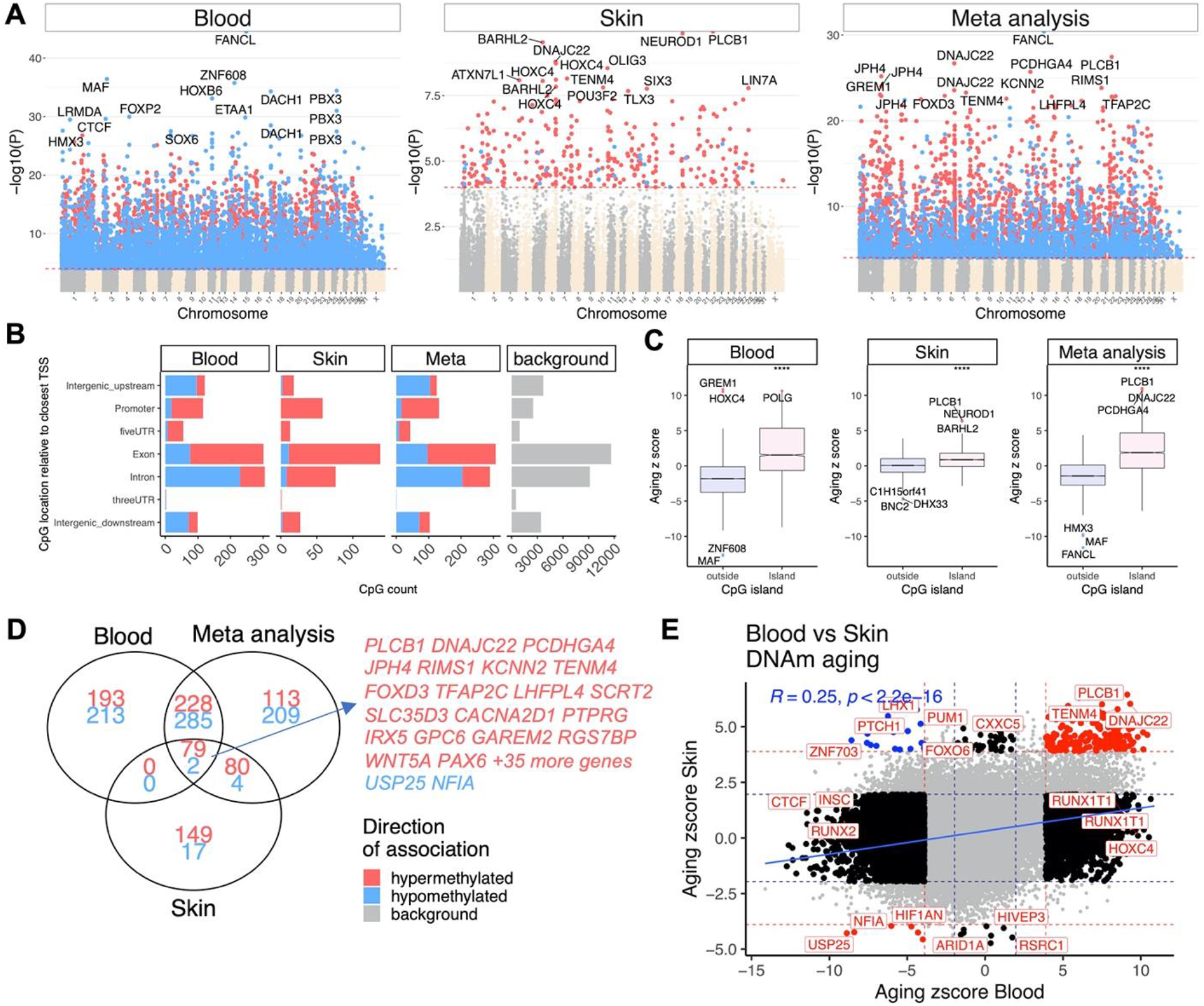
Epigenome wide association study (EWAS) of age in blood and skin of zebra. A) Manhattan plots of the EWAS of chronological age. Since a genome assembly was not available for zebra, the coordinates are estimated based on the alignment of mammalian array probes to EquCab3.0.100 (horse) genome. The direction of associations with p < 10^−4^ (red dotted line) is highlighted by red (hypermethylated) and blue (hypomethylated) colors. Top 15 CpGs was labeled by the neighboring genes. B) Location of top CpGs in each tissue relative to the closest transcriptional start site. Top CpGs were selected at p < 10^−4^ and further filtering based on z score of association with chronological age for up to 500 in a positive or negative direction. The number of selected CpGs: blood, 1000; skin, 331; meta-analysis, 1000. The grey color in the last panel represents the location of 3,1836 mammalian array probes mapped to EquCab3.0.100 genome. C) Box plot of a Z statistic resulting from a correlation test with age versus CpG island status. The median Z statistics are significantly different, p<10^−4^. D) Venn diagram of the top age related CpGs in blood and skin of zebra. E) Sector plot of DNA methylation aging in blood and liver of horse tissues. Red dotted line: p<10^−4^; blue dotted line: p>0.05; Red dots: shared CpGs; black dots: tissue specific changes; blue dots: CpGs whose age correlation differs between blood and skin tissue.

DNAm aging had a moderate positive correlation (r = 0.25), and with only 80 overlapped CpGs between blood and skin of zebras (**Figure 4D, E**). Some of these shared changes include hypermethylation in *PLCB1* exon, *RIMS1* exon, and hypomethylation in *NOVA2* intron and NFIA intron (Supplement 3, Figure S5). This moderate correlation could be due to a small number of skin samples rather than biological difference. Future studies should validate this observation by a larger sample size. As opposed to small overlap, age related CpGs in both blood and skin were related to development (e.g. nervous system), survival, and were enriched with polycomb repressor complex 2 (e.g. EED, SUZ12, PCR2) target genes (Supplement 3, Figure S6). This observation corroborates prior findings on DNAm aging in many other mammalian species [41].

## Discussion

To the best of our knowledge, this is the first study to present DNA methylation-based age estimators for any wild equid. The high accuracy of the epigenetic clocks (ECs) reflects that we used a custom array that profiled 36 thousand probes that were highly conserved across numerous mammalian species. These robust data allowed us to construct highly accurate epigenetic aging models for zebras. The best model to predict chronological age was the EC developed from blood samples, which predicted individual age +/− 6 months.

A challenge to obtaining a highly accurate biopsy-based clock may be the variability of tissue types within such a sample. Biopsy needles for remote darting are variable in width and depth. The needles used in this study were 1 mm wide and 25 mm long. Biopsy samples consist of three skin layers - the epidermis, the dermis and the hypodermis - and may even contain deeper tissues. These layers can vary in thickness across the body and among individuals and therefore may be present in different proportions across samples. In spite of the inherent difficulties of using biopsy samples and of the skewed age distribution (Supplement 3, Figure S4), our biopsy clock predicted age with an error of +/− 2 years.

The zebra blood EC accurately predicts the chronological ages of horses, Grevy’s zebras and Somali wild asses. This is not surprising; ECs developed for humans can be used to estimate age in chimps (Horvath, 2013), which are more distantly related (6-8 MYA, [37]), than caballine and non-caballine equids (4-4.5 MYA [38, 59]). Non-caballine species of equids are closer to plains zebras (1.28-1.75 MYA), than domestic horses so we would expect the blood EC developed for plains zebras to show even stronger correlations and lower error rates. In fact, chimpanzees and bonobos have a similar divergence time to those observed within the non-caballine equids [60] and align more closely to each other in DNAm age than with humans (Horvath 2013). In our analysis the plains zebra clock seemed to perform slightly better Grevy’s zebras and Somali wild asses on horses.

The epigenetic pacemaker models (EPMs) developed here reveal that epigenetic changes occur in a non-linear fashion throughout the lifespan of zebras. As has been found in humans and other species [31, 34], young zebras undergo faster epigenetic changes than adult zebras. We also observed that the variance in the estimates of epigenetic age is lower in young compared to old zebras. Increased variation in epigenetic age in adults is observed in humans and other species, and may be a consequence of lifetime accumulation of environmental and physiological factors on the epigenome [7, 14].

Genome wide increase in methylation levels has been observed in inbred plants [28, 30]. In salmon, inbreeding was associated with altered methylation of three genes [29], while thousands of alterations in methylation were detected in inbred chickens reversal of inbreeding depression has been achieved by demethylation of inbred lines in plants [30]. We found that EC-estimated epigenetic age is mildly accelerated in individuals with higher levels of inbreeding as measured by the genetic inbreeding coefficients F and F_ROH_. EPM-estimate age-acceleration was significantly associated only with the inbreeding coefficient F, even though F and FROH were strongly correlated. Although the high precision of ECs for predicting age can negatively impact their ability to detect age acceleration associated with biological variation [61, 62], our EC detected biological variation where the EPM did not. [63]. A lack of concordance between the two models suggests that the association between age-acceleration and inbreeding may not be robust.

The probes in this array were selected based on conservation in mammalian genomes, thus, our findings are expected to have high translatability into humans and other mammals. The high positive correlation with age observed in promoter CpGs in blood is indeed consistent with DNAm aging in humans and other species [64, 65]. The poor conservation across age effects may reflect the limited sample size in the biopsy samples (n=20) or it could point to biological differences between these two tissues. The GREAT analysis was based on the human genome, thus, the relevance of these pathways to zebra requires further validation. Regardless of this limitation, the enrichment analysis suggests a partially conserved aging biology between humans, zebra, and many other mammalian species.

The epigenetic aging models are expected to become valuable resources for conservation efforts in equids. Chronological ages predicted by ECs can be used to estimate reproductive potential and population viability. In known age populations, both ECs [27, 66–68] and EPMs [31–33] have the potential to identify causes of individual accelerated aging. These models can be particularly useful in combination with ecological or other genetic data [4]. Given the small number of sites required, an aging project in a wild population could be done relatively inexpensively using bisulfite sequencing or pyrosequencing rather than an array [69–71]. While epigenetic models based on blood are more accurate and less invasive than many other options for aging mammals, there are drawbacks in that animals must be immobilized to obtain the samples. Because biopsy samples can be obtained with minimal disruption [72], a highly reliable clock based on biopsy samples will be a worthwhile direction for future research. Since the methods for extracting genomic DNA from feces have been improved [73–75], it will be worthwhile to explore whether our epigenetic aging models can be adapted to this non-invasive source of DNA.

## Data accessibility

The data will be made publicly available as part of the data release from the Mammalian Methylation Consortium. Genome annotations of these CpGs can be found on Github https://github.com/shorvath/MammalianMethylationConsortium

## Author contributions

BL and SH conceived of the study. BL and GP analyzed data, and BL, GP and SH co-wrote the article. The remaining authors helped with the statistical analysis or the data generation. All authors reviewed and edited the article.

## Acknowledgements

This work was supported by the Paul G. Allen Frontiers Group (SH). GP was supported by the Science Without Borders program of the National Counsel of Technological and Scientific Development of Brazil. Sample collection was supported by National Geographic Grant 8941-11 (BL). We thank the following people for their assistance acquiring the plains zebra blood and tissue samples: Colleen O’Ryan, Stephen Mitchell, Mick D’Alton, Tom Turner, Boet le Roux, Evert Grobbelar, Ewald Groenewald, Basie and Coenraad Bezuidenhout, Linda Mason, Patricia Swanepoel, Fernando Rueda, Ross Cowlin, Melissa Stander, Hanna Lindstadt, Ansel Abels, J.P. Hugo, Cobus van Coller, Jannie du Plessis, and SANParks. Grevy’s zebra and Somali wild ass samples were graciously provided by White Oak Conservation.

## Supplemental Information for

### Supplementary Methods

#### RAD and Low Coverage Sequencing and Variant Calling

Library preparation and RAD sequencing was conducted as described in [1] using the enzymes SbFI and PacI. We aligned sequences to the horse genome EquCab3 [2] using BWA [3] and called genotypes using Freebayes [4] and Sentieon [5] to obtain a consensus set of 915476 variants. We removed indels, multiallelic and non-autosomal sites, loci genotyped in > 10% of individuals, and individuals genotyped at < 20% of loci. In addition, we kept loci with a mean depth between 5 and 75, and with a strand bias < 20, leaving 859122 SNPs.

Low coverage libraries were constructed from 200ng genomic DNA using Truseq Nano kit (Illumina), indexed with unique dual indices (Integrated DNA Technologies), and sequenced 24 libraries per lane on a HiSeqX platform (Illumina), generating 6.2±1.4G reads per sample. We aligned sequences to the horse genome EquCab3 [2] using BWA [3] and called pooled genotypes for the Quagga Project and two closely related source populations from Etosha National Park and Kruger National Park. Etosha and Hluhluwe National Parks are the exact locations where the Quagga Project founders were captured, but Kruger and Hluhluwe National Parks comprise a single population [1] using [4] and Sentieon [5]. In both datasets we removed indels, multiallelic sites, and those sites with < 0.1 MAF and depths more extreme than the median depth +/− 2IQR. We then filtered to select only SNPs common to both datasets. We used this set of 9100016 high confidence variants to generate allele frequencies at each site using Sentieon.

**Table S1.** Association between inbreeding and age acceleration as predicted by the plains zebra blood EC. N=66.

| **Variable** | **Coefficient** | **MAD** | **V value** | **P** |
| --- | --- | --- | --- | --- |
| *F* | 1.737 | 3.13 | 1685 | 0.0002 |
| *Length ROH > 1 MB* | 2.879 | 5.81 | 1664 | 0.0004 |
| *Length ROH > 3 MB* | 2.568 | 6.81 | 1536 | 0.006 |
| *Length ROH > 10 MB* | 2.724 | 10.01 | 1494 | 0.01 |

**Table S2.**
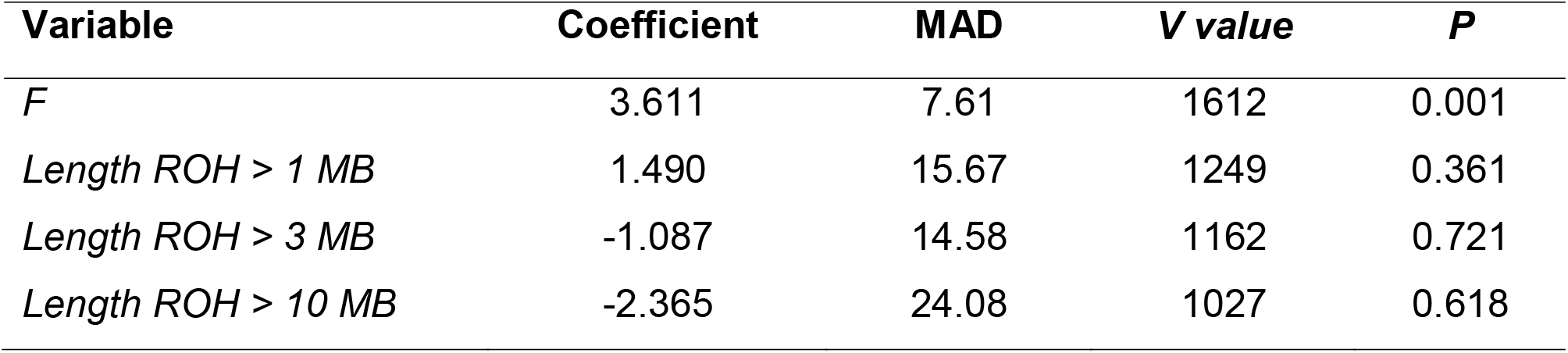
Association between inbreeding and age acceleration as predicted by the plains zebra blood EPM. N=66.

**Figure S1.**
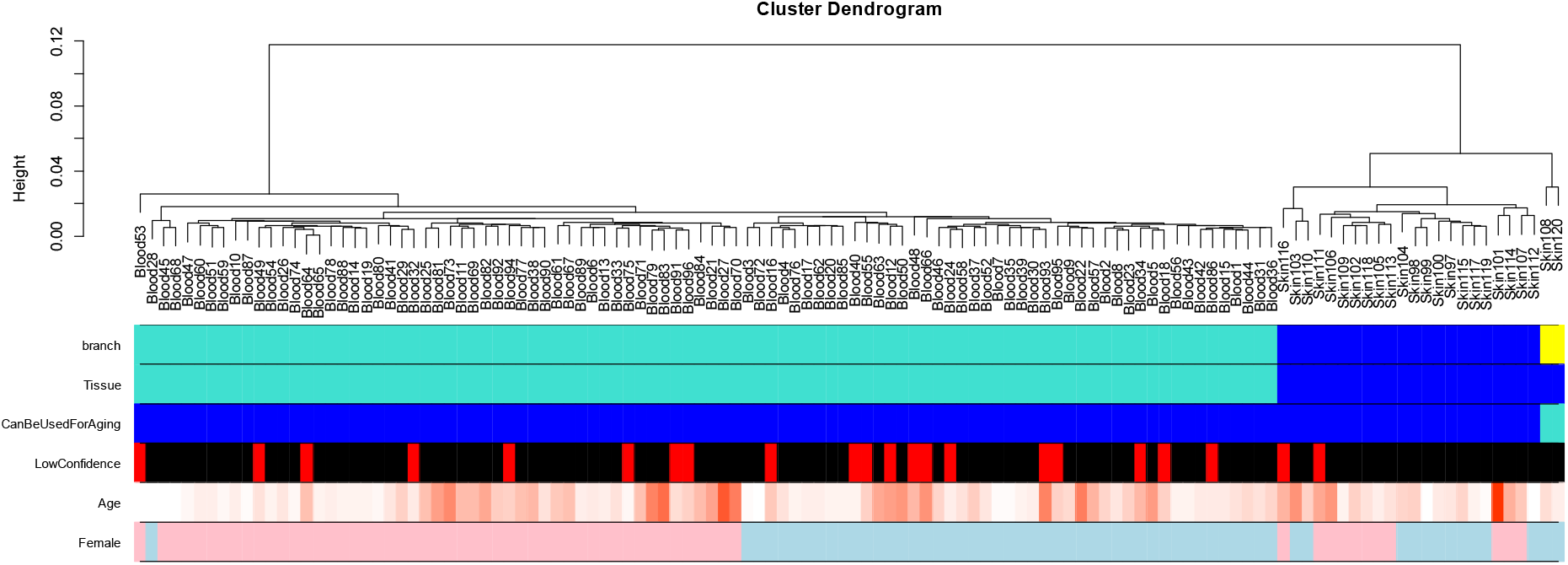
Unsupervised hierarchical clustering of blood and skin samples from zebras. Average linkage hierarchical clustering based on the interarray correlation coefficient (Pearson correlation). The relatively low height values (y-axis) indicate high inter array correlations and good quality. However, two skin samples cluster into a distinct yellow cluster (first color band). These putative outliers were removed from the analysis (third color band). Contrasting the first color band (based on cluster branches) with the second color band shows that the arrays cluster by tissue type (blue=skin). The fourth color band indicates wild animals whose ages were largely unknown (low confidence in the provided age estimate). These samples were omitted from the training set. The last color band indicates that the blood samples are grouped by sex.

**Figure S2.**
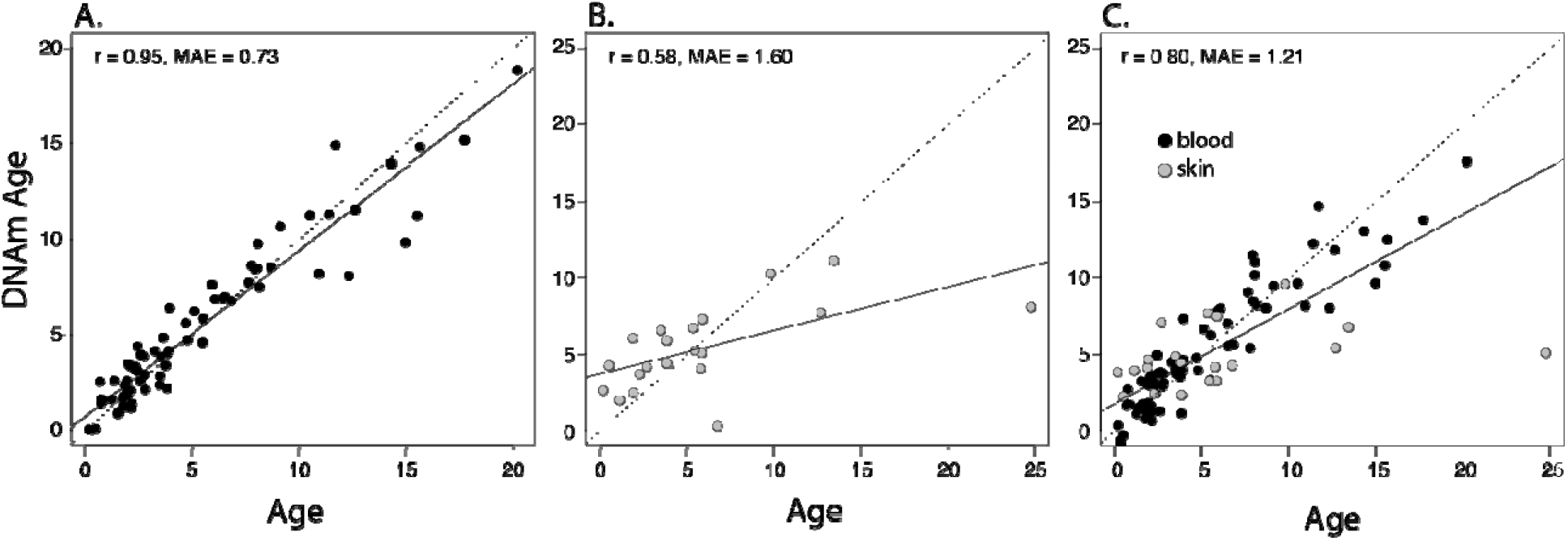
Epigenetic clocks using untransformed ages. We developed 3 epigenetic clocks for zebras based on untransformed ages: A) epigenetic clock for blood samples based in 215 CpG sites, B) epigenetic clock for skin samples based on 168 CpG sites, and C) combined tissue clock for both sample types based on 345 CpG sites. Leave-one-sample-out (LOO) estimate of DNA methylation age (y-axis, in units of years) versus chronological age (x-axis). The linear regression of epigenetic age is indicated by a solid line while (y=x) is depicted by a dashed line. Predictions based on 76 blood samples and 20 biopsy samples.

**Figure S3.**
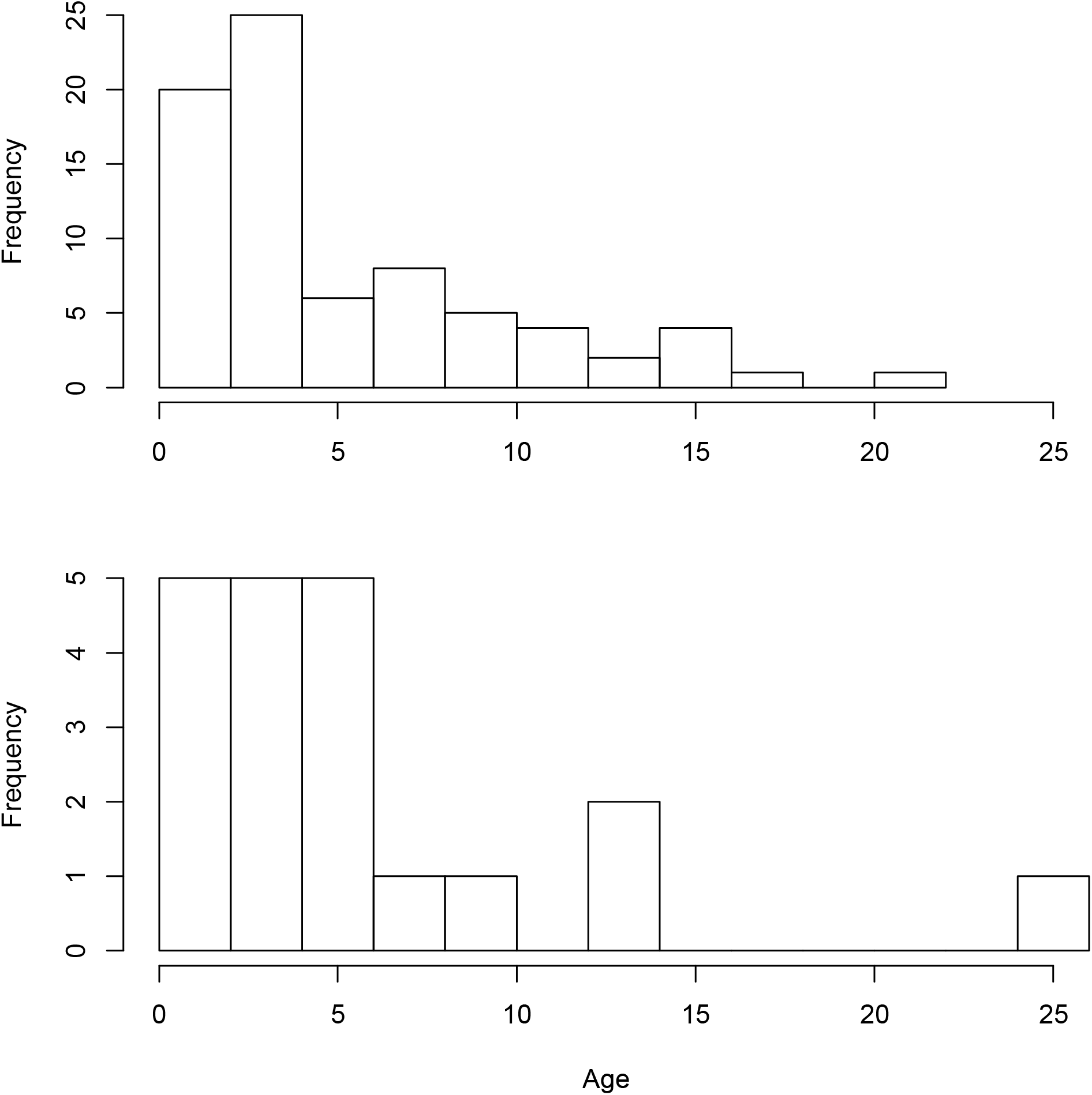
Association between inbreeding, F and ROH.

**Figure S4.**
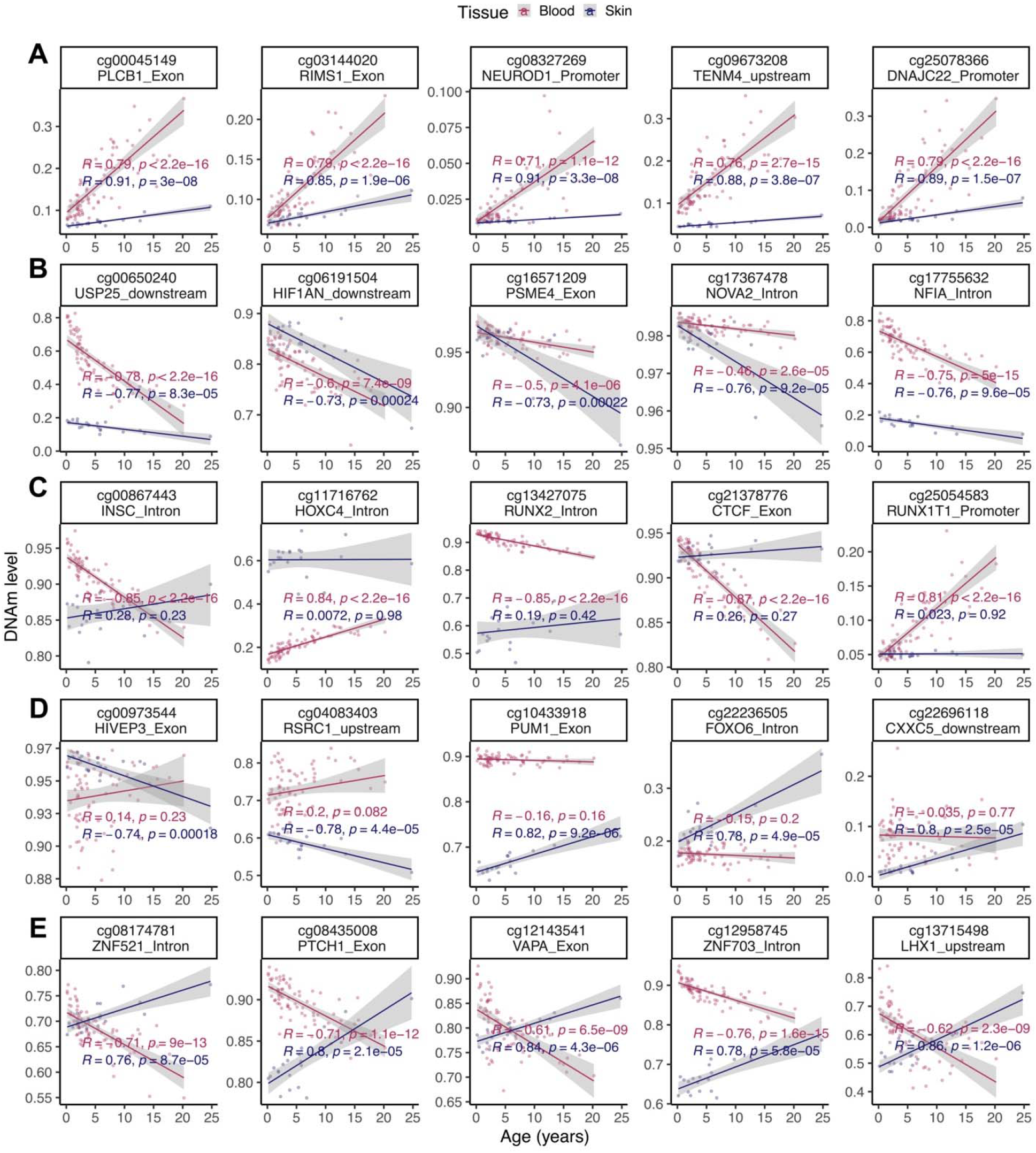
Age distribution of the samples. Top: blood samples, bottom: skin samples. **Scatter plots of age-related changes in selected CpGs blood and skin of zebras.** A) CpGs that are hypermethylated with age in both tissues. B) CpGs that are hypomethylated with age in both tissues. C) Examples of blood specific changes. D) Examples of skin specific changes. E) Selected CpGs with divergent aging pattern between skin and blood of zebras.

**Figure S5.**
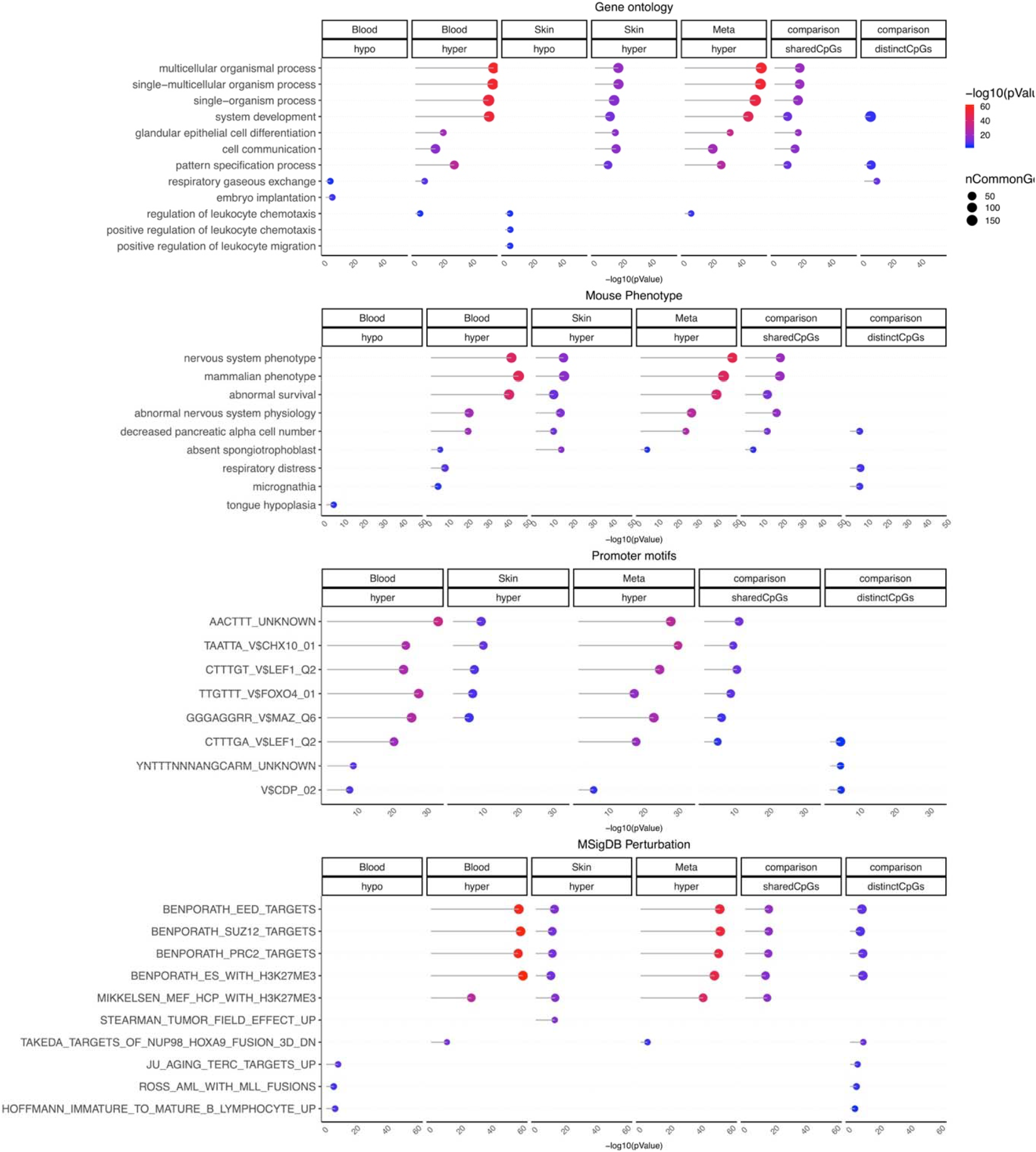
Gene set enrichment analysis of DNA methylation aging in zebra. The gene level enrichment was done using GREAT analysis [6] and human Hg19 background. Datasets: gene ontology, mouse phenotypes, promoter motifs, and MSigDB Perturbation. The results were filtered for significance at p < 10^−3^.

